# Logic-based modeling of biological networks with Netflux

**DOI:** 10.1101/2024.01.11.575227

**Authors:** Alexander P. Clark, Mukti Chowkwale, Alexander Paap, Stephen Dang, Jeffrey J. Saucerman

## Abstract

Molecular signaling networks drive a diverse range of cellular decisions, including whether to proliferate, how and when to die, and many processes in between. Such networks often connect hundreds of proteins, genes, and processes. Understanding these complex networks is aided by computational modeling, but these tools require extensive programming knowledge. In this article, we describe a user-friendly, programming-free network simulation tool called Netflux (https://github.com/saucermanlab/Netflux). Over the last decade, Netflux has been used to construct numerous predictive network models that have deepened our understanding of how complex biological networks make cell decisions. Here, we provide a Netflux tutorial that covers how to construct a network model and then simulate network responses to perturbations. Upon completion of this tutorial, you will be able to construct your own model in Netflux and simulate how perturbations to proteins and genes propagate through signaling and gene-regulatory networks.

## 1 Introduction

Biological signaling networks are responsible for a variety of cellular decisions, including growth, proliferation, disease progression, and death. These varied behaviors arise from interactions between hundreds of proteins, genes, and cellular processes. This complexity makes it difficult to understand mechanisms of cellular decisions through reductionist experimental approaches alone. Computational modeling addresses this challenge by integrating known interactions between system components into a framework; one can then use computer simulations to explore biological processes *in silico*, providing a means to rapidly generate testable predictions.

Understanding the signaling involved in biological processes can take decades of rigorous, reductionist cause-and-effect research. For example, a connection between high blood pressure and cardiac hypertrophy (often a precursor to heart failure) has been established for years, but until recently[1] (**Figure 1B**), the complex mechano-signaling pathways that drive this remodeling had not been integrated at a systems level. Over decades, individual studies have connected components (e.g., stretch activating integrin) and explained mechano-signaling pathways (e.g., a multi-protein pathway whereby stretch activates Ras) in cardiomyocytes. Collectively, these studies demonstrate the complex role of mechano-signaling in cardiac hypertrophy, but they do not provide a single mechanistic picture of stretch-induced hypertrophy.

Network models offer a means to integrate the high-quality reductionist building block studies mentioned above into a framework to study the complexity of biological signaling processes (**Figure 1A**). Building block studies identify the nature of interactions, like whether stretch increases or decreases the expression of calcium channels – it increases them. By mining many of these reductionist studies, one can create a network of interactions between species and construct an integrated picture of how individual interactions can lead to an emergent phenotype. For example, **Figure 1B** shows a mechano-signaling network for heart cells that was manually curated using data from over 170 studies [1]. This network includes mechanical Stretch as an input that stimulates several pathways consisting of receptors (AT1R, angiotensin type 1 receptor), ion channels (LTCC, L-type calcium channel), transcription factors (GATA4), sarcomeric proteins (bMHC, beta myosin heavy chain), and ions (Ca, Calcium). The interactions can be either activating or inhibiting. For example, the Stretch input signal increases (denoted as solid arrow) the expression of L-type calcium channels. In contrast protein kinase G 1 (PKG1) inhibits the expression of L-type calcium channels. The 125 activating or inhibitory interactions within this network give rise to emergent phenotypic characteristics, like Cell Area.

**Figure 1:**
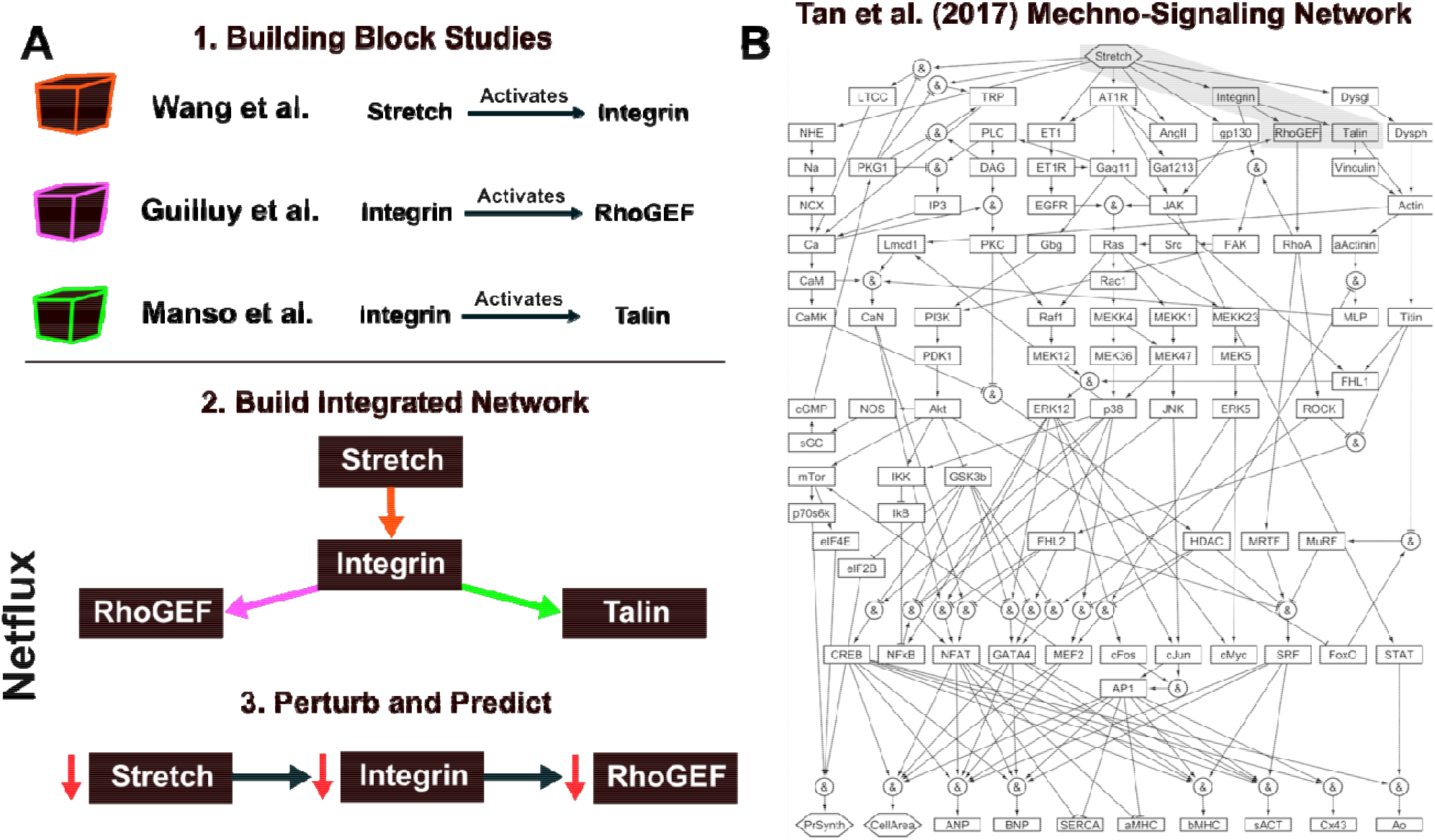
**A)** Individual reductionist studies provide connections between network components. These individual connections can be integrated into a network model. Subsequent perturbation studies can make novel predictions of previously untested genetic/protein perturbations – these come with a mechanistic explanation for the perturbation outcome. **B)** Network model from Tan et al. (2017) shows how complex networks can be constructed through manual curation. Subsequent simulations in this study provide novel insights into a biological process (in this case, cardiomyocyte mechano-signaling).

Coding these individual interactions into a cohesive network and then simulating genetic or pharmacologic perturbations allows us to uncover insights that are not apparent when studying components in isolation. Network modeling can prioritize pathways that work together or in tension to result in emergent phenomena, such as changes in Cell Area. By simulating these networks, researchers can explore how different pathways interact, identify key regulatory nodes, and predict system behavior under various conditions. For example, through systematic perturbation to the mechano-signaling network in [1], the authors identify mechanisms by which increased Stretch can increase Cell Area, a maladaptive change in heart cell physiology. Knowing the mechanism, they show how a recently approved combination heart failure drug called Entresto attenuates heart failure progression through distinct, yet synergistic, mechano-signaling pathways. This systems approach demonstrates how modeling and simulation can harness the complexity of molecular networks to identify biological mechanisms of disease progression and explain novel therapeutic approaches.

Despite decades of single-gene cause-effect research, the integration of these findings into comprehensive models has been hindered in part by the availability of user-friendly network modeling tools. As the amount of biological data continues to grow, and new methods/databases make it easy to find connections between molecules, there is a need for tools that facilitate the development and simulation of biological networks. In this manuscript, we provide a detailed introduction to Netflux, an intuitive tool for construction and simulation of biological network models that does not require programming.

Under the hood, Netflux utilizes a modeling approach termed logic-based differential equations, which is based on normalized Hill activation/inhibition functions [2]. Such an approach benefits from providing continuous variables to model gene and protein concentration, in contrast to traditional Boolean network modeling approaches with discrete variables. Netflux abstracts these technical considerations and allows users to construct networks and run simulations without any coding. If you’re interested in learning about the technical implementation of Netflux, more details on the underlying computational approach of logic-based differential equations can be found in Kraeutler et al. (2010, [2]). The tool has been used by scientists of various levels and technical proficiency to develop models of numerous biological networks that have led to meaningful insights and advances in our understanding of human disease [3–7]. Netflux has also been used as an educational tool at the undergraduate and high school levels to introduce the concepts of network biology.

By the end of this article, you will be able to use Netflux to create a biological network model and then predict how the network responds to perturbations in environmental conditions, proteins, and genes. We will also discuss advanced Netflux topics and provide published examples of how models built with Netflux have provided meaningful insights into biological processes.

## 2 Getting started with Netflux

### Installation

<u>Download</u> Netflux from GitHub (https://github.com/saucermanlab/Netflux). It can either be: option 1) installed as a desktop application, or option 2) opened and run directly from MATLAB.

### Loading a Model

<u>Open</u> the Netflux graphical user interface (GUI, **Figure** 2) by either clicking on the installed application (option 1) or running the “Netflux.m” file within MATLAB (option 2).

**Figure 2:**
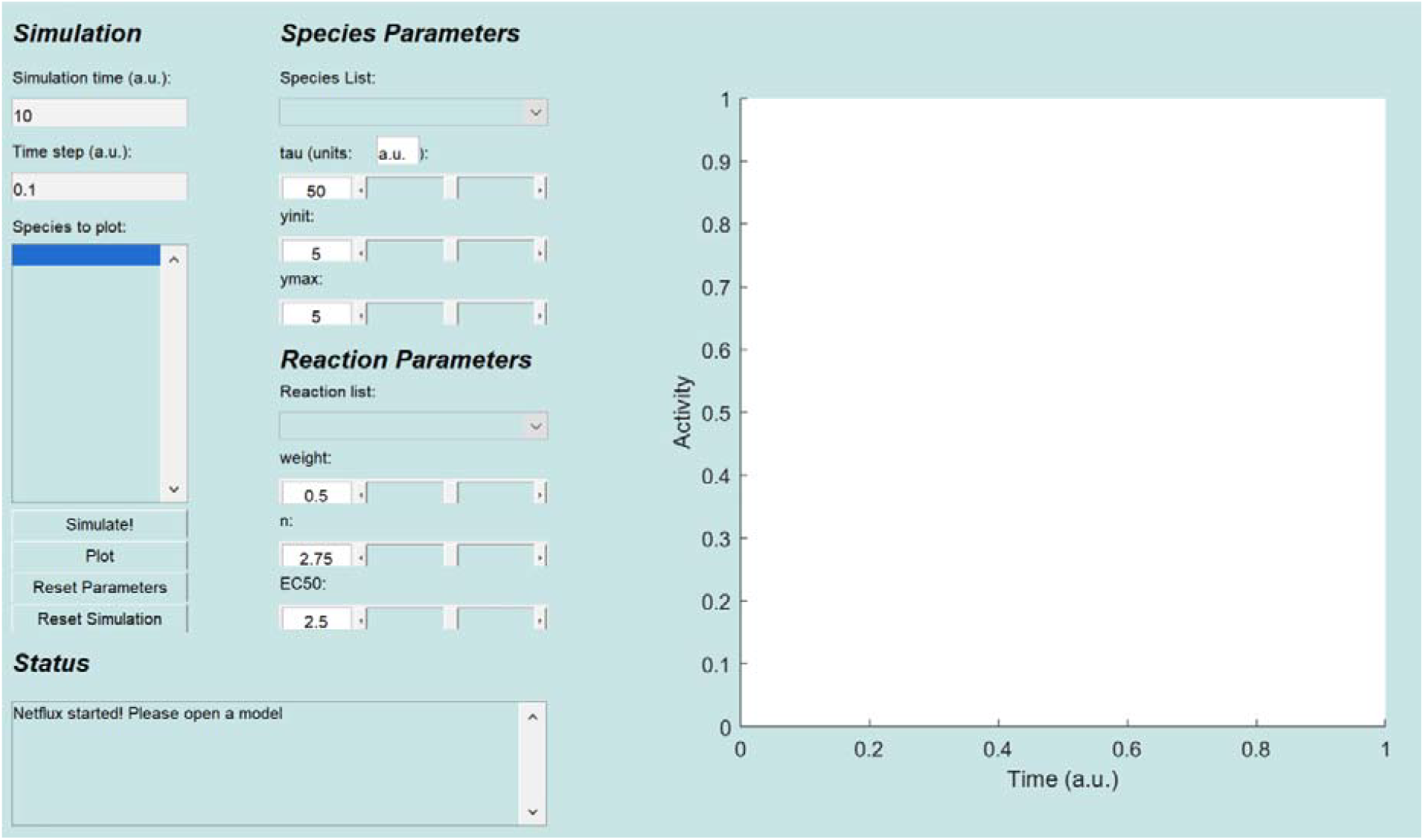
Netflux graphical user interface. The Netflux GUI includes four sections (left) and a plot (right) of species activity over time. The sections are: 1) ***Simulation*** includes information about the simulation time, and species that will be plotted in the panel on the right; 2) ***Status*** includes information on the state of the loaded model, and is important to check after loading models and running simulations; 3) ***Species Parameters*** includes information about the species initial value (yinit), maximum value (ymax), and how quickly it can change (tau); 4) ***Reaction Parameters*** includes information about a reaction’s strength (weight), the steepness of activation (n), and the half maximal effective concentration (EC50), which are collectively responsible for producing dynamic changes in species.

### Loading the ExampleNet Model

Netflux includes a repository of published network models in the “models” subfolder. We will first focus on the exampleNet model [2].

<u>Open</u> the “exampleNet.xlsx” file by selecting Open from the File tab within the Netflux GUI, and then navigating to models/exampleNet.xlsx.

In Netflux, we refer to network genes or proteins as *species*, and the activating or inhibiting interaction between species as a *reaction*. The ExampleNet model (**Figure 3A**) consists of five species (A, B, C, D, and E) and six reactions, including two input species (A and B) that feed into the network and can be turned on or off by turning on or off their respective input reactions.

- **(Input) Reaction 1:** Environmental stimulus that activates Species A.
- **(Input) Reaction 2:** Environmental stimulus that activates Species B.
- **Reaction 3:** Species A activates Species C.
- **Reaction 4**: Species B activates Species D.
- **Reaction 5**: Species A activates, and species B inhibits species E.
- **Reaction 6:** Species E activates species C.

**Figure 3:**
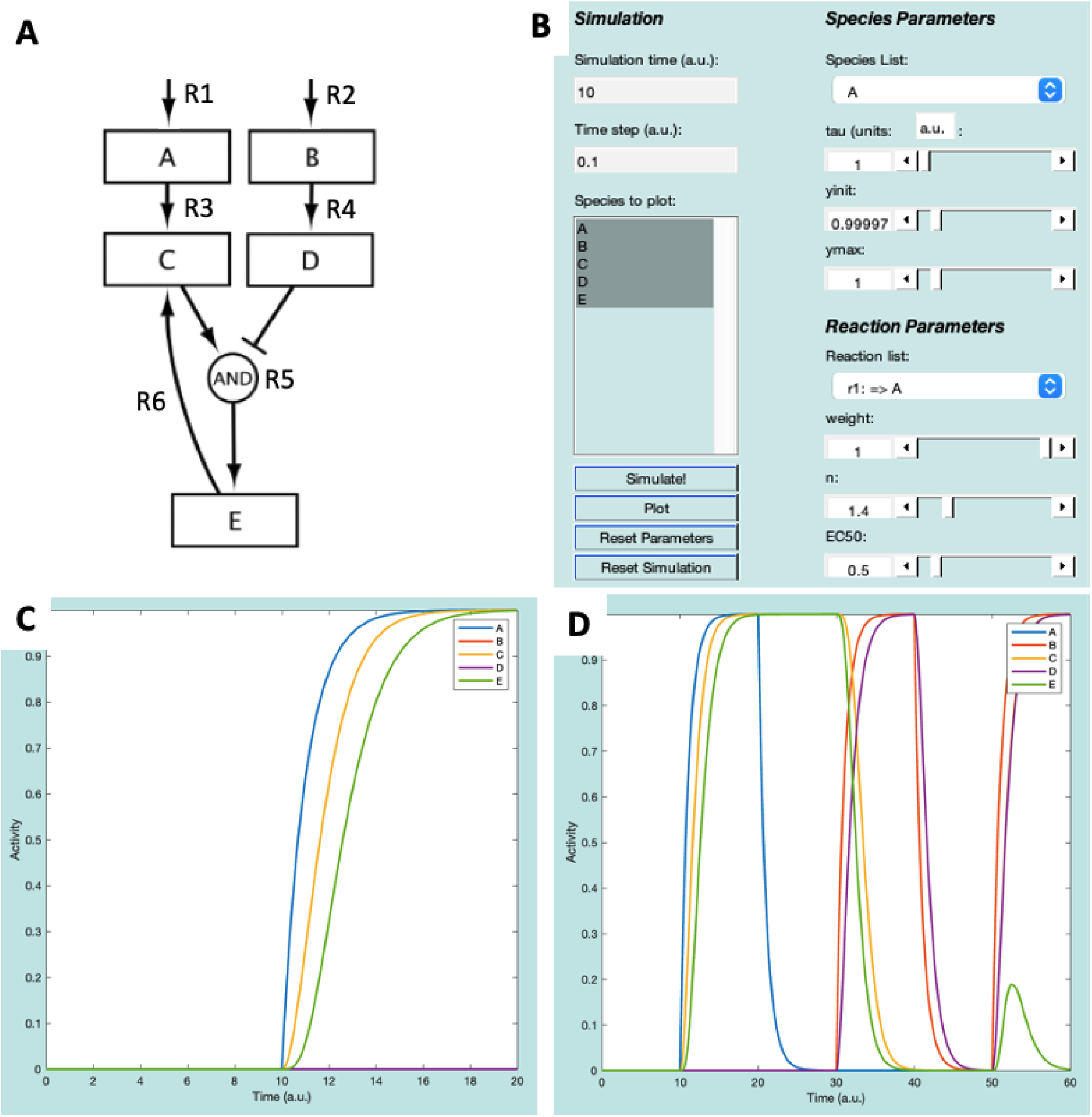
ExampleNet schematic (**A**) and GUI representation (**B**). (**C**) plot of species after Simulations 1 and 2. (**D**) Plot of species after all simulations.

Once loaded, all the network species are displayed in the “Species to plot” box of the Netflux GUI. Under ***Species Parameters***, <u>verify</u> that the values for time constants (tau=1), initial values (yinit=0), and maximum values (ymax=1) are the same for each species. Additionally, under ***Reaction Parameters*** (bottom right of the GUI), <u>verify</u> that the reaction weight (w=0 or 1), Hill coefficient (n=1.4), and half maximal effective concentration (EC50=0.5) are preset for each reaction.

### Simulating ExampleNet in Response to Environmental Stimuli

The model is ready to be run.

First, <u>select</u> all the species in the “Species to plot” box – this can be done by selecting the A, holding the “Shift” key, and then selecting E (**Figure** 3B). Follow the steps below and refer to the GUI in **Figure** 3B to run the model and plot its results:

1. <u>Press</u> the “Simulate!” button. The axes on the right side of the GUI should change, but none of the species should have an Activity above 0.
2. <u>Click</u> the “Reaction list” to see the reactions that are defined in ExampleNet. Change the reaction weight for “r1: => A” to 1, then click “Simulate!” again. Now, between timepoints 10 and 20, species A, C, and E should all increase to an Activity level of 1 (**Figure** 3C).
3. <u>Simulate</u> washout of A by changing the “r1: => A” reaction weight back to 0, and then clicking “Simulate!”.
4. <u>Simulate</u> stimulation with B by changing the “r2: => B” reaction weight to 1, and then clicking “Simulate!”.
5. <u>Simulate</u> washout of B by changing the “r2: => B” reaction weight back to 0, and then clicking “Simulate!”.
6. Finally, <u>simulate</u> simultaneous stimulation with both A and B by changing their input reaction weights to 1, and then clicking “Simulate!”.

Execution of these six steps will result in the plot displayed in **Figure** 3D. Before starting a new simulation, it is important to select “Reset Parameters” and “Reset Simulation”.

## 3 Build and simulate a new network model

This section will cover how to build a new model from scratch. The model used in this section is part of a recently elucidated gene regulatory network that is essential for proper heart formation [8], visualized in **Figure** 4A. In addition to model construction, we will simulate the knockout of a gene within this network and compare the results to experimental data linking dysregulation of this network to congenital heart defects.

**Figure 4:**
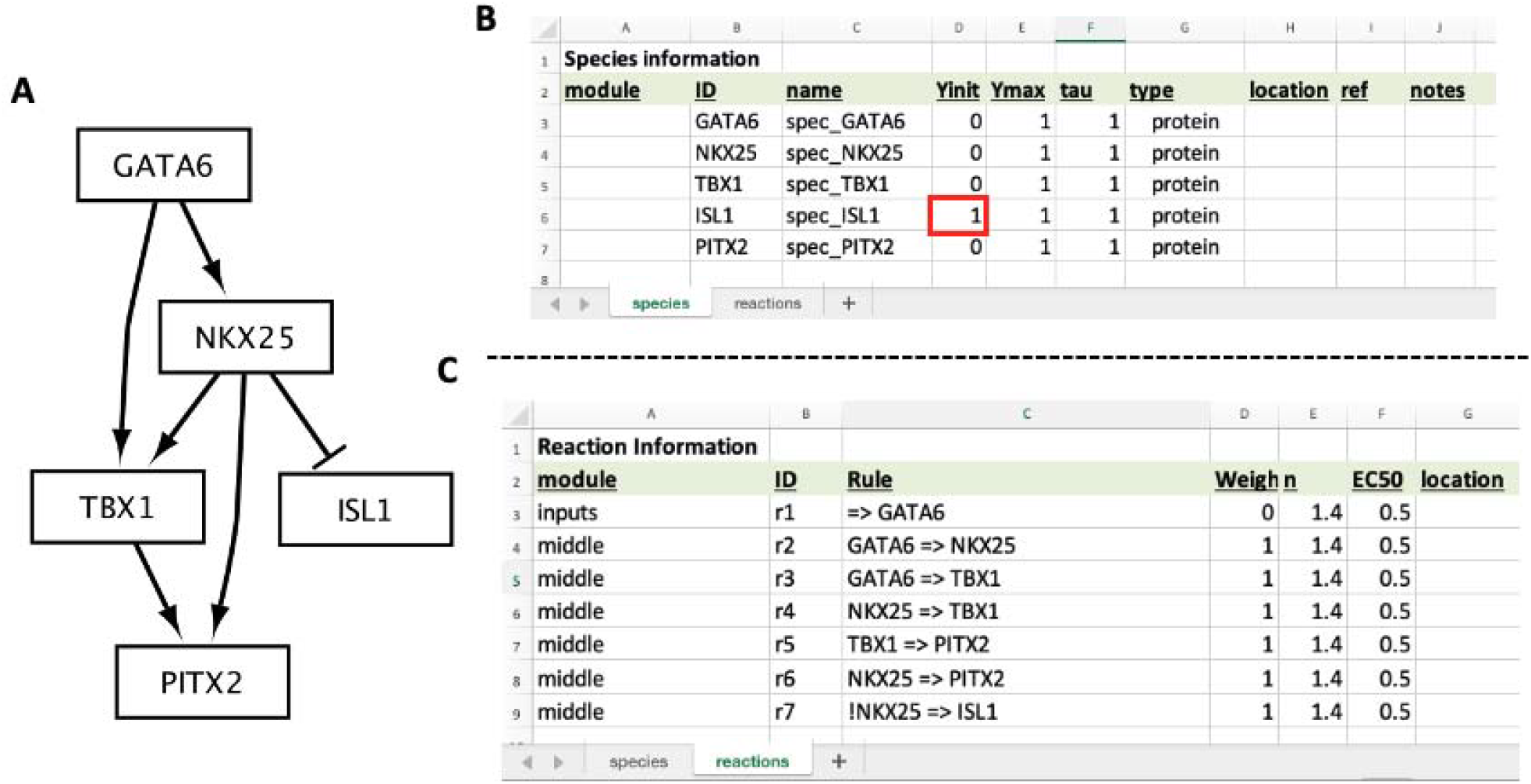
Model of a cardiac development gene regulatory network. (**A**) CardiacDevNet schematic. (**B**) Excel sheets for the CardiacDevNet Species and (**C**) Reaction information. Notice, the ISL1 Yinit value starts at 1 because it is inhibited by NKX25, which has a Yinit of 0 (**B**, red box)

### Creating the CardiacDevNet Model

First, <u>make a copy</u> of the “exampleNet.xlsx” model in Netflux/models/, and rename it “cardiacDevNet.xlsx”. Note, when building a new model, it is always advised to copy and then edit a previously developed model (e.g., “exampleNet.xlsx”).

### Adding the Model Species

This model includes five transcription factor species, with the following known actions (within this biological context):

- GATA6 – shown to bind an enhancer of NKX2-5 that is accessible in a special subset of cardiac progenitor cells found in the anterior second heart field.
- NKX2-5 – well-characterized protein that regulates several genes essential for proper heart formation.
- TBX1 – well-characterized protein that regulates several genes essential for proper heart formation.
- ISL1 – plays a role in forming the cardiac progenitor pool, but its repression is important for cells to continue differentiation towards adult cardiomyocytes.
- PITX2 – known to regulate formation of a region of the heart called the outflow tract.

<u>Open</u> “cardiacDevNet.xlsx” in excel or Google Sheets and <u>navigate</u> to the “species” tab. <u>Edit</u> the ID, name, and Yinit fields to match **Figure** 4A, B. Notice, the Yinit value for ISL1 should start at 1. This is because the subset of specialized cardiac progenitor cells (the anterior second heart field cells) starts with high levels of ISL1.

### Adding the Model Reactions

This model has the following seven reactions (**Figure** 4A, C), including one input reaction:

- **(Input) Reaction 1:** GATA6 is expressed. This is modeling the expression of GATA6 that is present within this specialized subset of cardiac progenitor cells.
- **Reaction 2:** GATA6 activates NKX2-5.
- **Reaction 3**: GATA6 activates TBX1.
- **Reaction 4**: NKX2-5 activates TBX1.
- **Reaction 5:** TBX1 activates PITX2.
- **Reaction 6:** NKX2-5 activates PITX2.
- **Reaction 7:** NKX2-5 inhibits the expression of ISL1. In Excel, an exclamation mark is used to denote that NKX2-5 inhibits ISL1 (**Figure** 4C, ID=r7).

<u>Navigate</u> to the “reactions” tab of “cardiacDevNet.xlsx”. <u>Edit</u> the module, ID, Rule, and Weight to match **Figure** 4C. For the “r1” reaction rule (“=> GATA6”), include a single quote before typing the equals character (=) to ensure excel properly interprets the coded interaction.

### Simulating Normal Development with CardiacDevNet

<u>Save</u> “cardiacDevNet.xlsx” in excel or export it from Google Sheets as a .xlsx file. Next, <u>open</u> Netflux by either clicking on the installed application or running the “Netflux.m” file within MATLAB and then <u>open</u> “cardiacDevNet.xlsx” (see Section 2 for details).

The following steps cover how to run a simulation of the normal cardiac development:

1. <u>Set</u> the simulation time to 2. This step improves visualization of results in this simulation. It is not always needed.
2. <u>Select</u> all species to plot. This is done by clicking on GATA6, holding the “Shift” key, and then selecting PITX2.
3. <u>Select</u> the “Simulate!” button. This should change the axes, but there should be no change in species Activity.
4. <u>Change</u> the simulation time to 8.
5. <u>Select</u> “r1: => GATA6” from the “Reaction list” dropdown and <u>change</u> the weight to 1.
6. <u>Select</u> the “Simulate!” button. The results should match the solid lines plotted in **Figure** 5B.

This simulation shows that NKX2-5, TBX1, and PITX2 increase while ISL1 decreases in response to an increase in GATA6 (**Figure** 5B, solid lines). These gene expression dynamics are important for anterior second heart field cells to progress towards mature outflow tract cardiomyocytes.

**Figure 5:**
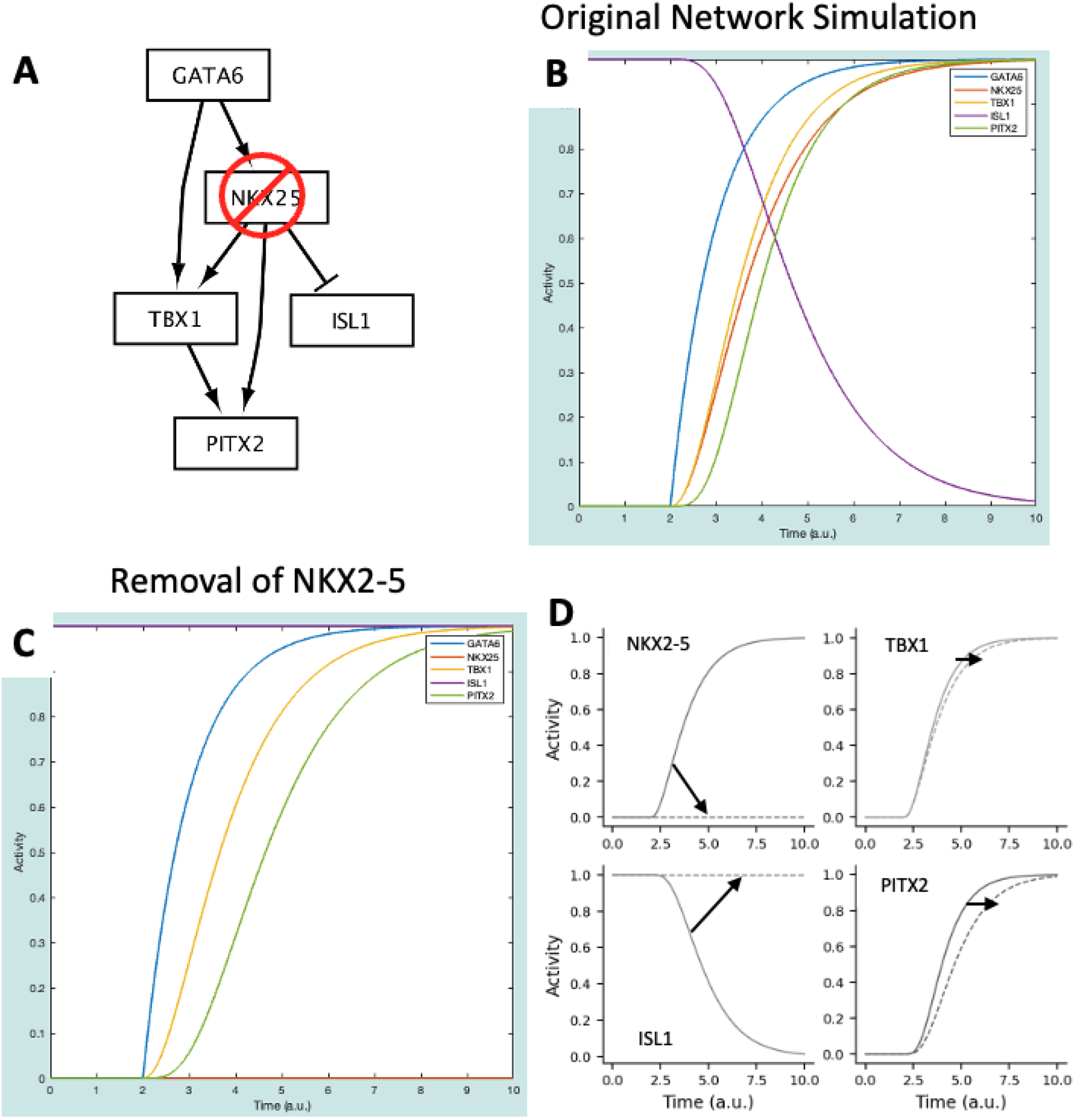
**(A)** CardiacDevNet NKX2-5 knockout schematic. **(B)**, CardiacDevNet simulation results at baseline. **(C)** CardiacDevNet simulation results after knockout of NKX2-5. **(D)** Change in simulation results at baseline (solid lines) and after NKX2-5 knockout (dashed lines) for the transcription factors: NKX2-5, TBX1, ISL1, and PITX2.

### Simulating a Gene Knockout withCardiacDevNet

In [8], authors reduce the expression of NKX2-5 – here, we simulate this as a knockout of the NKX2-5 gene. The next steps discuss how to conduct an *in silico* gene knockout study using the model (results displayed in **Figure** 5C):

1. <u>Select</u> “Reset Parameters” and “Reset Simulation”.
2. <u>Select</u> “NKX25” from the “Species List” dropdown and change the ymax to 0. This change simulates the knockout of NKX2-5.
3. <u>Set</u> the Simulation time to 2.
4. <u>Select</u> the “Simulate!” button.
5. <u>Select</u> “r1: => GATA6” from the “Reaction list” dropdown and change the weight to 1.
6. <u>Set</u> the Simulation time to 8.
7. <u>Select</u> the “Simulate!” button.

This simulation represents the conditions under which NKX2-5 is knocked out – this was done by removing NKX2-5 from the network (Figure **5**). Taken together, the results of the baseline and perturbation simulations predict the following effects of NKX2-5 knockout (**Figure** 5D), which largely agrees with the experiments in [8].

1. NKX2-5 is not expressed. This should be interpreted as a reduction in NKX2-5 expression compared to wildtype. In [8], while still present, NKX2-5 expression is significantly reduced by the removal of an enhancer region that, when bound by the GATA6 transcription factor, increases NKX2-5 expression.
2. ISL1 expression is not suppressed. This again agrees with the significant increase in ISL1 observed when NKX2-5 expression is reduced.
3. There is a lag in TBX1 and PITX2 expression. This can be interpreted as a reduction in expression, in agreement with [8], where TBX1 and PITX2 expression was reduced due to a reduction in NKX2-5.

## 4 Advanced methods and usage

Our simulations with CardiacDevNet illustrate the ease of developing a network model that accurately predicts the outcomes of real-world experiments. While small models such as CardiacDevNet are useful, Netflux has also been used extensively to construct much larger networks, with tens to 100s of proteins, genes, and reactions. Some such networks include: signaling networks for cardiomyocyte hypertrophy [3,9] and apoptosis [4], cardiac fibroblasts [6], macrophages [10], brain endothelial cells [7], mechano-signaling [1], virtual drug screening [11], cardiac differentiation [12], and smooth muscle cells [5]. As networks increase in size and complexity, there are a few additional considerations to be made.

### Network Visualization

Netflux provides a tool to export networks in a format that can be loaded into Cytoscape [13], a user-friendly software for network visualization. For example, Cytoscape was used to produce the visualizations in **Figures** 2-4.

### Citation Trackin

The Netflux-formatted .xlsx sheets include several columns within the “species” and “reactions” tabs that can be used to keep track of references and metadata. This becomes increasingly important as models grow large and incorporate insights from numerous independent studies.

### Model Validation

As noted above, novel network models often draw on many different independent, biologically targeted studies to define the network structure – this provides confidence in model definition. However, experimental validation is critical to build confidence in model predictions. Initial experimental validation is often performed using *in vitro* or *in vivo* perturbation experiments from the literature that were not used to develop the model. Subsequently, the model can be used to design novel experiments that test model predictions under previously untested conditions.

### Model Exporting

Netflux also provides functionality for exporting models to MATLAB or Python, making it possible to run more sophisticated analyses and conduct large-scale, systematic perturbation studies.

## 5 Conclusion

Netflux is an accessible tool for constructing and simulating models of biological networks, requiring no programming experience. In this article, we covered the core features of Netflux, including model construction, simulation, perturbation, and were able to produce *in silico* results in agreement with experimental data. While the models used in this article are small compared to networks from many published studies, the concepts remain the same. This article (along with information in the GitHub repository) provides all the information needed to construct your own model and simulate real experiments. As you embark on building network models, we encourage you to rigorously validate your model throughout construction. Well-validated models developed in Netflux can provide both conceptual understanding of biological networks and specific mechanistic predictions of how your network may respond to new experimental perturbations.

## Acknowledgements

We thank the members of the Saucerman lab for their feedback and contributions to Netflux, with special thanks to Taylor Eggertsen and Bryce Murillo for their feedback on this manuscript. Research reported in this publication was supported by the National Heart, Lung and Blood Institute of the National Institutes of Health under Award Number T32 HL007284.

